# Memory systems integration in sleep complements rapid systems consolidation in wakefulness

**DOI:** 10.1101/2023.03.09.531360

**Authors:** Svenja Brodt, Monika Schönauer, Anna Seewald, Jonas Beck, Michael Erb, Klaus Scheffler, Steffen Gais

**Affiliations:** Institute of Medical Psychology and Behavioral Neurobiology, University of Tübingen, Tübingen, Germany; Max-Planck-Institute for Biological Cybernetics, Tübingen, Germany; Department of Psychology, Neuropsychology, University of Freiburg, Freiburg, Germany; Department of Psychology, University of Marburg, Marburg, Germany; Swiss Sleep House Bern, Department of Neurology, Inselspital, Bern, Switzerland; Biomedical Magnetic Resonance, Universitätsklinikum Tübingen, Tübingen, Germany

## Abstract

Sleep benefits memory performance by fostering systems consolidation, a process that embeds memories into neocortical networks and renders them independent of the hippocampus. Recent evidence shows that memory rehearsal during wakefulness likewise initiates systems consolidation and rapidly engenders neocortical engrams. Here, we investigate the effect of sleep-dependent consolidation for memories that have undergone rapid systems consolidation during wakefulness. After sleep compared to wakefulness, we find better memory retention and higher functional brain activity during memory retrieval in the medial parietal cortex, which hosts memory representations after rehearsal, and in the striatum and thalamus. Increased striatal and thalamic contributions were correlated with higher retrieval performance. Furthermore, all three regions decreased their functional connectivity to the hippocampus specifically after sleep. These findings show that besides continuing of systems consolidation initiated during wakefulness, sleep also acts to integrate different memory systems. Thus, rehearsal-induced and sleep-dependent consolidation seem to be complementary in nature.

## Introduction

From many studies we know that sleep can benefit memory consolidation (Diekelmann & Born, 2010; Rasch & Born, 2013). To achieve this on a neural level sleep concurrently strengthens and weakens select dendritic spines and affects excitation-inhibition balance (Miyamoto et al., 2021; Niethard et al., 2021; Yang et al., 2014). While sleep’s role in memory is well studied on the behavioral level and at the microscale, evidence on how sleep operates on a systems level currently remains scarce.

For declarative memory, our memory for facts and events, an active process of systems consolidation has been proposed as one of the main mechanisms underlying the beneficial effect of sleep (Klinzing et al., 2019). Systems consolidation describes the process over which a freshly encoded hippocampus-dependent memory trace evolves to a stable representation in neocortical networks. It is achieved via transient reactivations of the distributed trace coordinated by hippocampal signaling (Frankland & Bontempi, 2005; McClelland et al., 1995). This so-called “replay” preferentially occurs during offline periods such as sleep (Bendor & Wilson, 2012; Buzsaki, 2015; Ji & Wilson, 2007; Wilson & McNaughton, 1994). Indeed, a night of sleep has been shown to shift the contribution of hippocampal and neocortical systems to memory retrieval on the short-term as well as on the long-term (Gais et al., 2007; Takashima et al., 2009). Whereas higher-order associative cortical regions like the medial prefrontal cortex and the posterior parietal cortex displayed an increase in memory-related functional activation and showed higher intracortical functional connectivity after sleep, hippocampal activity as well as hippocampo-neocortical functional connectivity decreased, indicating increased reliance of memory on intracortical networks.

Traditionally, systems consolidation was considered a primarily time-dependent transfer of mnemonic information from the hippocampus to the neocortex, yet recent evidence has changed our view towards a more flexible interaction of two complementary systems depending on many additional factors other than time (Brodt & Gais, 2020). New methods to tag and optogenetically manipulate engram cells have shown that neocortical engrams are formed from the outset of learning and that there are conditions under which they can mature rapidly within only one day (de Sousa et al., 2019; Kitamura et al., 2017). One factor promoting early neocortical memory formation across species are stimulus repetitions (Antony et al., 2017). In animals, increased training intensity has been shown to shorten the timeframe during which fear memories become hippocampus-independent (Lehmann et al., 2009; Pedraza et al., 2016). Likewise, human neuroimaging studies combining functional and microstructural measures have revealed that repeated encoding and retrieval lead to the emergence of a neocortical memory representation within a single experimental session for different kinds of declarative learning material (Brodt et al., 2018; Brodt et al., 2016; Himmer et al., 2019). Concurrently, memory-related hippocampal activation decreases, as does functional connectivity between the hippocampus and engram-hosting neocortical areas, indicating growing independence from hippocampal signaling.

These findings are strikingly similar to the results obtained when comparing memory maturation across a period of sleep, suggesting that external reactivation through rehearsal mirrors the natural process of internal, hippocampus-driven replay for systems consolidation (Antony et al., 2017). Consequently, the possibility that systems consolidation in the sense of changing hippocampo-neocortical contributions can be achieved during online learning raises the question whether and how sleep further contributes to the consolidation of these early neocortical traces. There is evidence indicating that sleep is needed to stabilize the hippocampal disengagement effected by repeated encoding and retrieval (Himmer et al., 2019), but it is currently unknown whether sleep also continues to shape neocortical representations. Interestingly, sleep can subserve memory consolidation not only by modulating the contribution of different nodes within a domain-specific mnemonic system (like hippocampus and neocortex within the declarative memory system), but also by the way how entire memory systems interact (Albouy et al., 2013; Cousins et al., 2016; Durrant et al., 2013; Orban et al., 2006). For example, in motor sequence learning, a night of sleep can change the co-activation of the involved declarative and procedural memory systems from a competitive to a cooperative mode (Albouy et al., 2013).

Therefore, in this study, we investigated the effect of sleep on the consolidation of repeatedly encoded and retrieved object-location associations previously shown to be represented in the medial posterior parietal cortex (mPPC) after only four rounds of rehearsal (Brodt et al., 2018). We were specifically interested in potential sleep-related changes in memory systems contribution within and beyond the reorganization already achieved through stimulus repetitions. Therefore, in a day-wake night-sleep design, two groups of participants underwent repeated runs of a memory task resembling the game ‘pairs’ in two sessions spaced 12 h apart while we recorded functional brain activity with fMRI (Fig. 1).

**Fig. 1.**
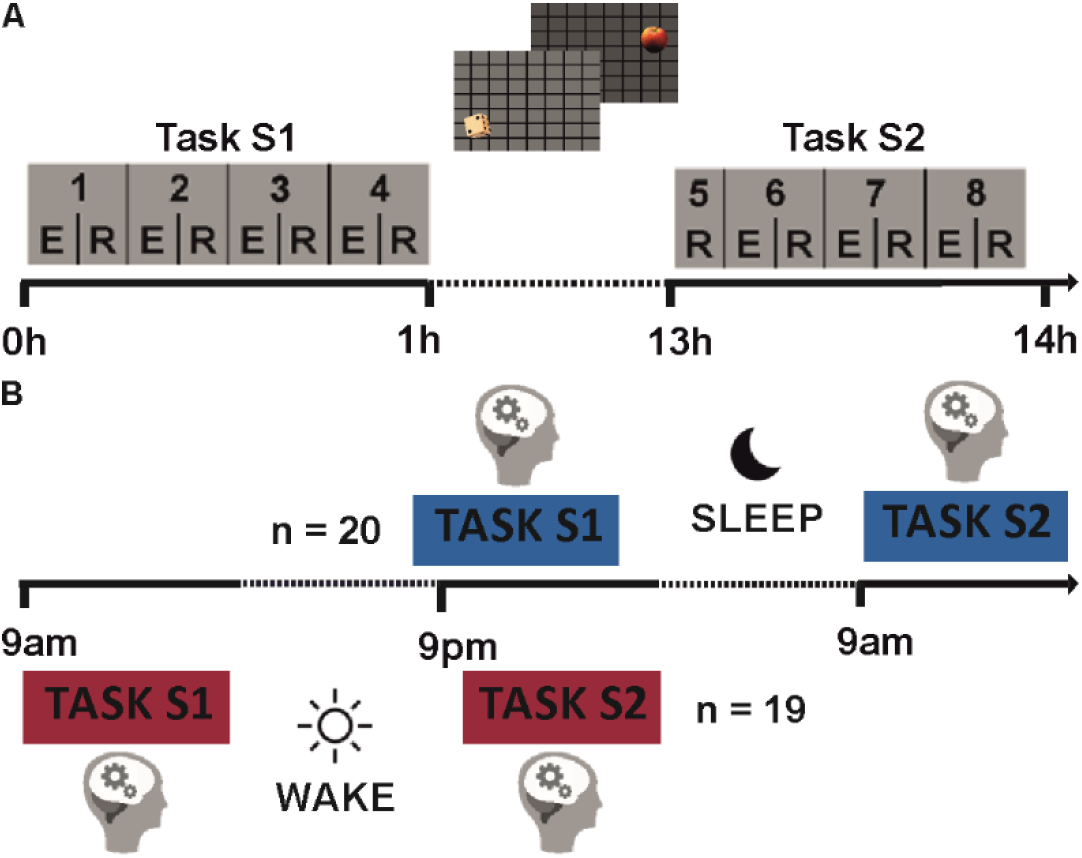
Experimental design. (**A**) Participants learned 40 associations between two items and their respective locations on a 2D grid over a total of 8 repeated encoding-retrieval runs distributed across two task sessions, which were spaced 12 h apart. (**B**) Participants in the sleep group attended the first session (S1) in the evening, then went home for a night’s sleep monitored via mobile polysomnography and returned for the second session (S2) the following morning. The wake group completed S1 in the morning, stayed awake during the day and returned in the evening for S2.

## Results

### Sleep preserves memory retrieval performance

To establish whether sleep benefits memory retention at the behavioral level, we examined patterns of retrieval performance in the sleep and wake groups across repeated learning and the retention interval. In both groups, memory performance increased rapidly over the first session, with a mean correct retrieval rate of 67.6±21.4% [M±SD] in the last retrieval run of the first session in both groups. During the second session, performance continued to increase and then plateaued at a ceiling level of 86.0±16.1% [M±SD] in the final retrieval run. However, while performance in the wake group decreased significantly from the last run of S1 (run 4, ‘pre’) to the first run of S2 (run 5, ‘post’), performance in the sleep group remained more stable across the 12-h delay (Fig. 2A; interaction effect: *F*_1,37_=6.24, *p*=.017, η^2^=.144; wake group: *t*_18_=6.10, *p*<.001; sleep group: *t*_19_=2.05, *p*=.054). Interestingly, this difference in retrieval performance persists as a trend until the end of the session, following three additional encoding-retrieval runs (Fig 2B; R8 *t*_37_=1.83, *p*=.075). Furthermore, there was a trend that the time participants spent asleep during the retention interval correlated with better memory retention (Fig. 2C; *r*_20_=0.43 *p*=0.056). Conversely, sleep did not affect overall reaction times (wake pre: 6.76±1.62 sec, wake post: 6.96±1.41, sleep pre: 6.60±1.47, sleep post: 7.13±2.12, [M±SD]; interaction effect: *F*_1,37_=0.64, *p*=.43, η^2^=. 144).

**Fig. 2.**
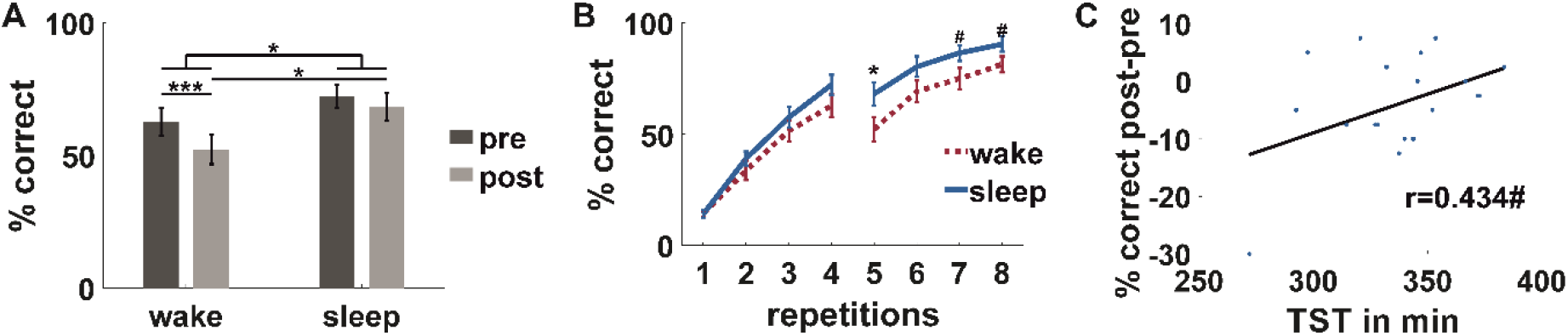
Sleep benefits memory retention. (**A**) The wake and the sleep group differ in memory retention. Across a retention interval of 12 h, retrieval performance from the last run of S1 (pre, dark grey) to the first run of S2 (post, light grey) decreased significantly more in the wake group than in the sleep group. (**B**) A trend for better memory in the sleep (blue solid line) compared to the wake group (red dotted line) is preserved across additional encoding-retrieval runs until the end of S2. (**C**) Participants with longer total sleep time (TST) tend to maintain memories better across the retention interval. Black line represents regression, blue dots represent individual data points. Data are M±SEM. # *p*<.08, * *p*<.05, *** *p*<.001.

### Sleep benefits memory retention by enhancing mnemonic activity in a posterior cortical and subcortical memory network

We next examined how sleep affects functional brain activity during mnemonic processing. Analogous to the behavioral analysis, sleep effects on BOLD responses during retrieval were assessed via interaction contrasts with the factors group (wake/sleep) and session (pre/post). The main analysis excluded time-of-day effects (Muto et al., 2016; Orban et al., 2020) via residual analysis. It showed significant sleep effects in clusters in the precuneus, putamen and thalamus (Fig. 3A and Table S1, *p*_SVC_<.05). Post-hoc analyses of the corresponding mean beta values of the significant voxels as well as the corresponding anatomically defined ROIs showed that this interaction was driven by a concurrent decrease in functional activity in the wake group and an increase in the sleep group across the retention interval in all three areas (Fig. 3B and Table S3A, *p*_SVC_<.05, for anatomical ROIs see Fig S1 and Table S3C). An additional analysis of the data unadjusted for time-of-day effects confirmed these findings (Fig. 3A+C, Table S2 and Table S3B, *p*_SVC_<.05). Moreover, regions displaying significant time-of-day effects do not overlap with those showing sleep effects (Table S1 and Table S4), excluding time-of-day as a principal explanatory factor for memory-related effects.

**Fig. 3.**
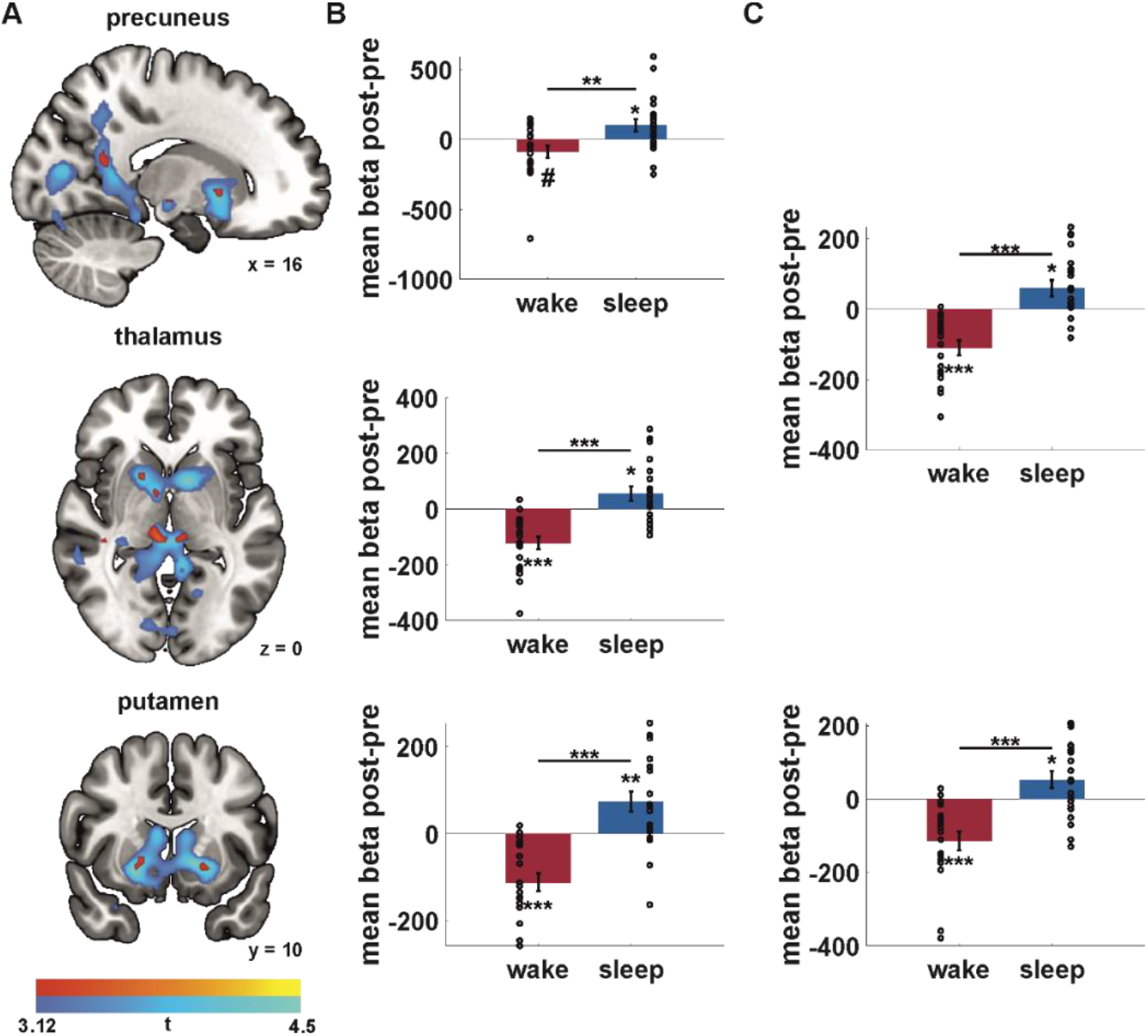
Sleep increases retrieval activity in a cortico-subcortical memory network. (**A**) Clusters located in the precuneus, thalamus and putamen show a significant modulation of retrieval activity by sleep in a conservative residuals analysis excluding effects of time spent awake at the time of data recording (red, interaction group*time, Table S1). Clusters are overlaid on results from the same analysis on the original unadjusted data (blue, Table S2). (**B**+**C**) Mean beta values of the interaction effect. Red, wake; blue, sleep; dots represent individual data points. See Table S3A-B. For the residuals (**B**, Table S3A) as well as the original data (**C**, Table S3B), the interaction is driven by a concurrent decrease in retrieval activity in the wake group and an increase in the sleep group. For corresponding analysis over all voxels in the anatomically defined ROIs see Table S3C. Clusters exhibit significant peak-level effects for the interaction analysis at small-volume corrected *p*_SVC_<.05, exceed 10 voxels and are displayed at *p*_uncorr_<.001. All beta values are corrected for baseline activation levels. Data are M±SEM. # *p*<.065, * *p*<.05, ** *p*<.01, *** *p*<.001.

Changes in functional activity in thalamus and putamen were correlated with memory retention across time (Fig. 4B-C): Participants with better memory retention also displayed a stronger increase in the retrieval-related response of putamen and thalamus across the retention interval (thalamus r_39_=.392 *p*=.014; putamen r_39_=.389 *p*=.014). No such relationship emerged between functional activity and change in reaction time (thalamus r_39_=-.125 *p* = .449; putamen r_39_=-.130 *p* = .430), excluding reaction time as mediating factor.

**Fig. 4.**
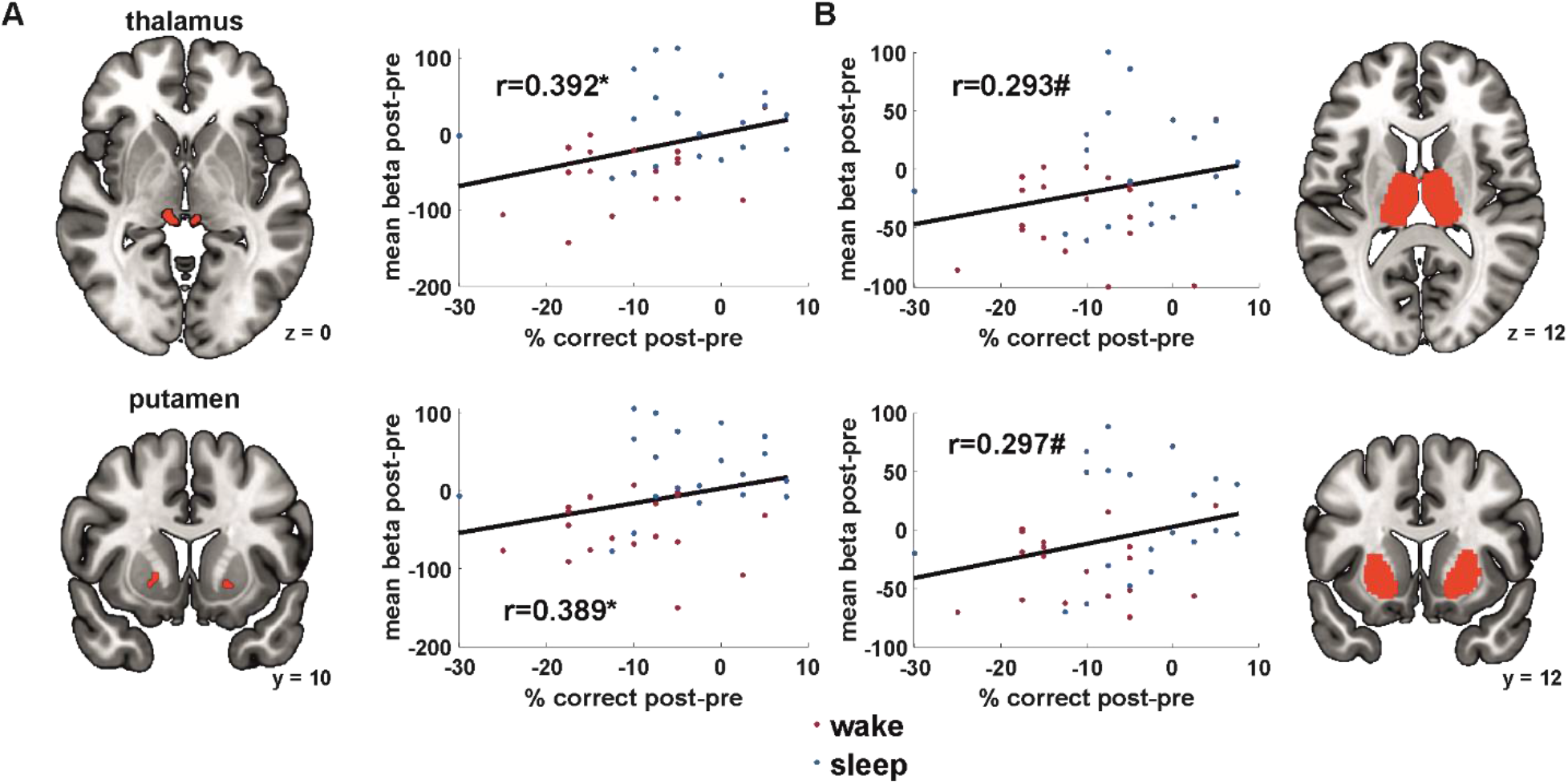
Sleep-dependent increases in thalamic and striatal retrieval activity benefit memory retention. In the thalamus and putamen, the strength of retrieval activity increase across the retention interval was correlated with memory retention performance across participants. Dots represent individual data points; red, wake; blue, sleep; black line, linear regression. (**A**) Mean beta values of the significant clusters from the residual analysis. (**B**) Mean beta values across all anatomically defined ROI voxels. Note that there is no difference in correlation between the wake and the sleep group (Fisher’s Z-Test, max. Z=0.33, min. p=.739). Clusters exhibit significant peak-level effects at small-volume corrected *p*_SVC_<.05, exceed 10 voxels and are displayed at *p*_uncorr_<.001. All beta values are corrected for baseline activation levels. #*p*<.071, \**p*<.05.

### Sleep-dependent increases in precuneus, thalamus, and putamen activity coincide with decreases in connectivity with the hippocampus

Increased consolidation of memories in neocortical regions is supposed to replace the need for hippocampal contributions to memory retrieval. We therefore investigated functional connectivity between regions showing sleep-dependent consolidation and the hippocampus in a whole-brain psycho-physiological interaction (PPI) analyses, which used anatomically defined seed regions. Interestingly, all three regions displaying robust sleep-related increases in retrieval activity also showed sleep-related changes in functional connectivity to the hippocampus (Fig. 5A and Table S5, *p*_SVC_<.05). Analysis of mean cluster beta values yielded a significant decrease in connectivity in the sleep group and an increase in the wake group driving the interaction effect (Fig. 5B and Table S6A). These effects hold not only for the significant cluster, but can also be seen when including all voxels within the whole anatomically defined hippocampus (Fig S2 and Table S6B).

**Fig. 5.**
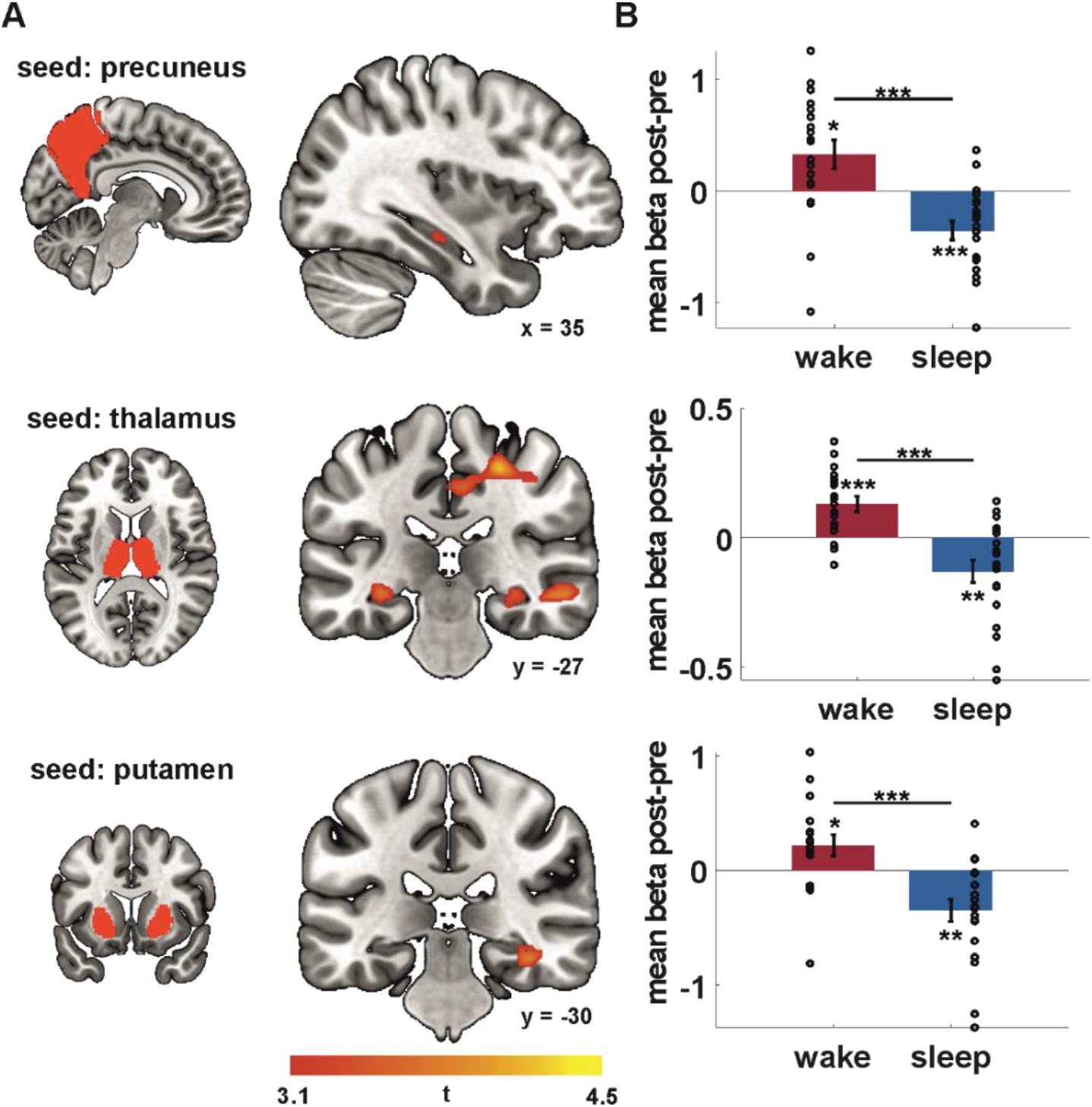
The precuneus, thalamus and putamen show sleep-dependent decreases in memory-related functional connectivity with the hippocampus. (**A**) Whole-brain PPI analyses with the anatomically defined precuneus, putamen and thalamus as a seed region indicate a significant sleep effect on connectivity with the hippocampus (interaction group*time, Table S5). (**B**) Investigation of the mean beta values of the significant clusters shows a decrease in connectivity with the hippocampus in the sleep group and an increase in the wake group across the retention interval (Table S6A). For corresponding analysis over all voxels in the anatomically defined hippocampus ROI see Fig S2 and Table S6B. Data are M±SEM. Dots represent individual data points. Hippocampal clusters exhibit significant peak-level effects at small-volume corrected *p*_SVC_<.05. Depicted clusters exceed 10 voxels and are displayed at *p*_uncorr_<.001. All beta values are corrected for baseline activation levels. * *p*<.05, ** *p*<.01, *** *p*<.001.

## Discussion

In this study, we investigated the influence of sleep on the consolidation of memories that have been repeatedly studied and retrieved. These memories, have previously been shown to have a neocortical representation (Brodt et al., 2018). We found that a night of sleep compared to a day awake stabilizes memory performance on the behavioral level and increases functional brain activity during retrieval in the precuneus, thalamus and striatum. Activity in the thalamus and striatum was correlated with better memory retention. Moreover, sleep reduced connectivity between all three regions and the hippocampus, indicating that memory processing in the posterior cortical and subcortical memory systems becomes more independent from hippocampal signaling over sleep.

In a previous analysis of this dataset (Brodt et al., 2018) and in other studies (Brodt et al., 2016; Himmer et al., 2019), we could already show that after completion of a single learning session, there is a memory engram in the precuneus, which manifests in microstructural modifications visible in diffusion-weighted MRI, and a distinct shift in retrieval-related functional brain activity towards the mPPC and away from the hippocampus. Because those findings are already indicative of a systems consolidation process, it was a central question of the present study whether sleep can play a further role in consolidation, going beyond the changes induced by previous repeated encoding and retrieval during wakefulness.

It has been argued that memory reactivation during repeated retrieval might lead to rapid neocortical integration and hippocampal independence, potentially forestalling further systems consolidation across sleep (Antony et al., 2017). Indeed, memories that were learned via fast mapping, which is a mnemonic strategy that fosters rapid neocortical integration (Coutanche & Thompson-Schill, 2014; Merhav et al., 2015; Sharon et al., 2011), do not benefit from sleep (Himmer et al., 2017). The behavioral data of the present study however show that sleep facilitates retrieval of rehearsed memories, in line with previous reports (Abel et al., 2019; Antony & Paller, 2018; Bäuml et al., 2014). This beneficial effect had a medium to large effect size and tended to persist even across multiple additional post-retention rehearsal opportunities. Sleep can therefore be assumed to have a beneficial effect even on memories that have already undergone a systemic, structural change in storage location.

The imaging results show that sleep serves to stabilize and enhance the pattern of activity initiated during rehearsal. Functional activity in the precuneus, the same region that has previously been shown to develop a memory engram of the task (Brodt et al., 2018), further increases over a night of sleep while it decreases over a daytime period of equal length without sleep, even when controlling for potentially confounding effects of time spent awake. Concurrently, functional connectivity with the hippocampus decreases. This pattern is similar to what has previously been found for the influence of sleep on declarative material (Gais et al., 2007; Takashima et al., 2009), and it is indicative of a continued systems consolidation process, which is initiated through rehearsal during learning and continues in sleep (Brodt et al., 2018; Brodt et al., 2016; Himmer et al., 2019; Schott et al., 2019). Interestingly, word list rehearsal leads to similar mPPC functional activity changes and a sleep-dependent stabilization of rehearsal-initiated hippocampal disengagement (Himmer et al., 2019).

These findings suggest that the continuation of neocortical consolidation, initiated prior to sleep, is one important contribution of sleep within declarative memory. Together with the reduction of connectivity to the hippocampus, both aspects thought to be relevant to systems memory consolidation – the shift in memory systems utilization and independence of the hippocampus – occur during sleep even when rapid systems consolidation has already occurred during wakefulness. The independence of the neocortex from the hippocampus, manifest in the reduced connectivity, seems to be a distinctive feature of sleep, as this was not yet seen at the end of the encoding session.

Besides a continuation of systems consolidation within the hippocampo-parietal declarative memory system, our data revealed sleep-related increases in retrieval activity in the putamen and the thalamus that were correlated with better memory retention. From a traditional memory perspective, these findings are surprising, because hippocampal and striatal networks have largely been viewed as segregated systems supporting different memory domains, i.e., declarative and procedural memory, respectively (Knowlton et al., 1996). However, they corroborate recent evidence that suggests that memory systems can also function interactively, especially after sleep.

Typically, the striatum is known for its participation in acquisition, consolidation, and retrieval of motor memories, especially in sequential motor tasks (Doyon et al., 2009). Offline gains correlate with increased striatal activity during and after sleep (Debas et al., 2010; Peigneux et al., 2003; Vahdat et al., 2017). However, there is also evidence for a contribution of the putamen to certain episodic memory tasks, particularly at later learning stages (Sadeh et al., 2011; Scimeca & Badre, 2012; Wang & Voss, 2014). Interactions between the medial temporal lobe and striatal memory systems have been mostly investigated in memory paradigms entailing both procedural and declarative components, such as motor sequence learning or perceptual statistical learning. Here, changes in functional connectivity between these systems have been observed across consolidation windows of 24 h and there are indications that these changes might depend on sleep (Albouy et al., 2008; Durrant et al., 2013). Furthermore, in spatial navigation tasks that can be solved with either hippocampus- or striatum-dependent strategies (Doeller et al., 2008), sleep has been shown to affect functional connectivity as well as behavioral strategy usage (Noack et al., 2021; Orban et al., 2006). The nature of hippocampus-striatum interactions seems to depend strongly on the specific task at hand (Freedberg et al., 2020), with some studies reporting decreased functional connectivity after sleep (Durrant et al., 2013; Orban et al., 2006), while others emphasize an integrative function (Albouy et al., 2013; Noack et al., 2021).

In our data, the sleep-related decrease in hippocampo-striatal connectivity emerged in parallel with a progression of systems consolidation, resulting in increased neocortical and striatal contributions to memory retrieval independent of hippocampal signaling. Together with the positive correlation between putamen activity increase and memory retention, these results support the idea of a cooperative relationship between neocortical and striatal memory nodes that emerges across sleep-dependent offline consolidation. The specific function of the striatum remains open for now. It might support successful retrieval by adaptively selecting from parietally stored memory representations based on their learned value and admitting them into working memory for correct response selection (Scimeca & Badre, 2012). Another possibility is that it integrates motor routines into the explicit spatial response (Knowlton et al., 1996).

The contribution of thalamic nuclei to mnemonic processing, especially during the encoding and retrieval stages, is well-established (Aggleton & Brown, 1999; Carlesimo et al., 2015; Van der Werf et al., 2003). While the thalamus has mostly been ascribed auxiliary functions for memory such as directing attention and controlling executive resources (Van der Werf et al., 2003), it has recently been shown that thalamic input to sensory regions is required for associative learning because it conveys mnemonic information about the behavioral relevance of an incoming stimulus (Pardi et al., 2020). Moreover, thalamo-cortical connectivity during consolidation periods predicts long-term memory, showing that the thalamus is not only involved in information transmission during stimulus presentation but also in consolidation-related processing (Wagner et al., 2019). Likewise, remote retrieval displays a functional reorganization of thalamo-hippocampal-cortical circuits towards a more hub-like role of the thalamus (Wheeler et al., 2013), emphasizing its importance for memory systems consolidation.

In the present study, we find a behaviorally relevant and sleep-dependent upregulation of the ventral midline regions of the thalamus during memory retrieval and a simultaneous decrease in functional connectivity between the thalamus and the hippocampus. This finding indicates that consolidation of memories during sleep restructures the memory trace in a way that involves multiple processing systems. It is in line with evidence for a differential involvement of the thalamus in the retrieval of recent vs. remote memories. While thalamic lesions leave memory acquisition and recent memory retrieval intact, remote memory expression is impaired (Loureiro et al., 2012).

The thalamus contributes to the information transfer between the hippocampus and higher-order neocortical areas especially for repeated stimuli by mediating hippocampal -cortical functional connectivity (Reagh et al., 2017). In humans, systems consolidation has previously already been associated with decreased functional connectivity between the thalamus and the hippocampus (Thielen et al., 2015). We show that this decoupling mainly occurs over sleep. Recently, evidence has accumulated that thalamo-cortical sleep spindles might play a coordinating role for information flow in hippocampal-neocortical circuits (Latchoumane et al., 2017; Yang et al., 2019). Particularly, the sleep-dependent emergence of experience-induced visual cortex plasticity critically depends on intact connections between thalamus and early visual areas (Durkin et al., 2017), emphasizing the role of the thalamus in promoting neocortical plasticity during sleep.

Together, our data complement the existing literature of a role of the thalamus in memory consolidation in general and sleep-dependent systems consolidation in particular by showing a behaviorally relevant contribution of sleep-dependent changes in thalamic activity and connectivity. Importantly, in this study, thalamic activity specifically benefitted systems consolidation of memories dependent on the mPPC, indicating a role of the thalamus beyond its known influence on early sensory areas and the medial prefrontal cortex.

In this study, we investigated the effect of a night of sleep on declarative memories that have already undergone rapid neocortical consolidation through repeated rehearsal (Antony et al., 2017; Brodt et al., 2018). We found evidence for a continued systems consolidation process during sleep that strengthened neocortical memory traces in the mPPC and rendered them less dependent on the hippocampus. At the same time, we observed a sleep-dependent enhancement of memory retrieval associated with an important node of the procedural memory system, the putamen, and with an important coordinating hub between memory systems, the thalamus. The behavioral relevance of their contributions emphasizes the importance of these two regions in the consolidation of declarative memories during sleep and might hint at a specific integrative function of sleep that cannot be accomplished solely by rehearsal. However, future studies employing more extensive rehearsal paradigms will be able to further test this idea and potentially reconcile our findings of continued systems consolidation during sleep with the literature showing a lack of sleep effects on purely neocortical memories (Himmer et al., 2017; van de Ven et al., 2016).

## Methods

### Participants

41 healthy, right-handed subjects (22 female, 19 male; mean age, 24.16±2.65 years [M±SD]) participated in this study, 20 in the sleep group, 21 in the wake group. Data from two participants in the wake group were excluded due to bad memory retrieval performance (more than 2 standard deviations below the population mean). Experimental procedures were approved by the Ethics Committee of the Medical Faculty of the Eberhard Karls Universität Tübingen, participants gave written informed consent and were compensated with 12€/h.

### Memory task

During the experiment, participants had to encode and retrieve item-location associations. The task resembled the game ‘pairs’: participants were shown pairs of ‘cards’, each depicting a unique object of everyday use in a specific location on an 8×5 grid on the computer screen. Each of the 40 cards was part of two different pairings, once as the first item and once as the second item of a pair, resulting in a total of 40 associations to be learned. During encoding, card placeholders (‘back sides’) for all 40 positions were shown. Then, paired items were presented sequentially for 2s each, separated by a short delay jittered between 0.4–0.6s. One encoding run consisted of presenting all pairs sequentially in random order, separated by an inter-trial interval jittered between 1.55–1.95s. During cued recall, participants were presented with the first item of a pair and had to indicate the location of the second item. All 40 pairs were again presented in random order. Participants were instructed to first think carefully about the second item’s location and only afterwards navigate a cursor to the respective position. They responded via two custom-built hand-shaped 5-button boxes (Max-Planck-Institute for Biological Cybernetics, Tübingen, Germany). The four principal navigation directions were mapped to the buttons for the index and middle fingers (up/down – r/l-index, right/left – r/l-middle), with button presses effecting a cursor jump to the next location on the grid in the respective direction. Participants confirmed their choice with the right thumb. All participants completed 4 encoding and 4 retrieval runs in the first task session, and a further retrieval followed by 3 additional encoding and retrieval runs in the second task session. Encoding as well as retrieval runs were interrupted every 60–90s by a baseline task lasting between 6–9s, asking them to fixate a white cross on black background that was moving between the grid positions at the same speed as item presentation during encoding. The task was programmed with Cogent 2000 (v1.33, Wellcome Trust Centre for Neuroimaging, London, UK) in MATLAB R2015b (Mathworks, Sherbom, MA).

### Procedure

Aspects of the data in this manuscript were published previously (Brodt et al., 2018). That paper focused on rapid brain plasticity induced by the first task session, the focus of the present analysis is the independent effect of sleep or wakefulness between the first and second session.

Participants were scanned in two sessions spaced 12h apart and randomly assigned to one of two groups. The sleep group completed the first session in the evening (mean task start time: 20:13±0:50h [M±SD]) and the second session in the following morning (9:33±0:54h), whereas the wake group had the first session in the morning (09:07±0:57h) and the second session in the evening of the same day (22:27±1:07h). During the first session, participants first completed a training phase for all constituents of the memory task (encoding, retrieval, baseline) outside of the MR scanner, consisting of a smaller grid version (4×3 locations) with different stimuli than the ones used in the actual task. This was done to ensure correct understanding of the instructions (especially the importance of the sequence information) and to practice the use of the button box. The first session consisted of two MR scanning blocks. During the first block, a series of control, anatomical, and diffusion-weighted scans were acquired (see MRI Data Acquisition and Analysis). Immediately afterwards, the task fMRI acquisition started. During this first task session, participants completed a total of four encoding-retrieval runs with a short break after two runs. Mean task duration was 50.93±4.50min [M±SD]. The task was projected to a screen at the back of the scanner bore, which the participants could see via a mirror attached to the head coil. Afterwards, while waiting for the second MR scanning block, participants stayed in the lab, were given a light meal and were allowed to read a recreational book or watch movies, but were expressly forbidden to study.

The second MR scanning block occurred approximately 3h after the start of the first block and consisted of the same series of control, anatomical, and diffusion-weighted scans as the first block. Afterwards, participants were taken out of the scanner and sent home. While the sleep group was equipped with a mobile EEG for polysomnography during the night, participants in the wake group were given a wristband actimeter (ActiSleep Monitor, ActiGraph, Pensacola, FL) to validate that they stayed awake during the day. Participants were not allowed to study, do any sports, or consume alcohol during the interval.

Upon returning to the lab for the second experimental session, participants completed a questionnaire about their activities during the interval and then proceeded to the third MR scanning block. The MR sequence protocol was similar to the first block. This time, the task fMRI session started with a retrieval-only run to assess retention performance, followed by three more encoding-retrieval runs. The second task session occurred approximately 13h (800±11 min [M±SD]) after the beginning of the first task session and the mean task duration was 41.81±4.17min [M±SD]. There was no difference between the groups in either task duration or retention interval between the sessions (task duration S1: *t*_37_=-0.63, *p*=0.534; task duration S2: *t*_37_=1.27, *p*=0.213; retention interval: *t*_37_=-0.06, *p*=0.954).

### Behavioral data and performance measures

All analyses in relation to behavioral performance reported in this article are based on the percentage of correctly retrieved locations of the second item in response to the presentation of the first item of each pair, separately for each of the retrieval runs. Statistical testing of performance measures relied on univariate ANOVAs or in the case of comparisons at single time points on two-sample t-tests. All tests were two-tailed with an α-level of 0.05.

For correlation analyses with fMRI data and sleep EEG parameters the difference between the retrieval runs directly before and after the retention interval was taken into account, i.e. the first run of session 2 (run 5, ’post’) minus the last run of session 1 (run 4, ’pre’).

### Sleep data

Sleep of the participants in the sleep group was monitored via a mobile EEG (Somnoscreen plus, SOMNOmedics GmbH, Randersacker, Germany), enabling them to sleep at home in their usual environment. Eight Ag-AgCl electrodes were attached immediately after the end of the second MR scanning block and were positioned at C3 and C4 of the international 10-20 system to record EEG, above the musculus masseter to record EMG and diagonally transposed left and right of the eyes to record EOG. Reference and ground electrodes were located on the right mastoid and the right collar bone, respectively. After checking electrode impedance, participants were sent home. Once lying in bed and ready to fall asleep, they pressed a button on the recording device, providing the ‘lights off’ time point. Similarly, the next morning after waking up a second button press indicated that the participants were awake and getting up.

Data was preprocessed with BrainVision Analyzer (v2.2, Brain Products GmbH, Gilching, Germany). Data were segmented according to the manual ‘lights off’ and ‘wake up’ markers. All channels were passed through a 50 Hz notch filter, EEG channels were band-pass filtered between 0.5 and 30 Hz, EMG channels were high-pass filtered above 25 Hz and EOG channels were low-pass filtered below 10 Hz with a Butterworth zero-phase filter of 48 dB/oct. Sleep scoring was done manually in 30 s epochs according to Rechtschaffen and Kales (1968) with inhouse software operating on MATLAB (R2019a, Mathworks, Sherbom, MA) by two independent raters. Sleep characteristics are reported in Table 1. Total sleep time (TST) and sleep delay (DEL) calculated from the end of task session S1 to sleep onset in minutes were considered for correlation analysis with memory performance.

**Table 1.** Sleep architecture for the retention interval night in the sleep group. Data are displayed in minutes [M±SD] and percent of TST. TST = total sleep time; S1-4 = sleep stage 1-4; REM = rapid eye movement sleep; DEL = sleep delay from end of the task. Note that in the 12-h interval, participants were required to travel to their homes to sleep, explaining the long sleep delay.

|  | minutes | % TST |
| --- | --- | --- |
| TST | 337±28 | - |
| S1 | 26±11 | 7.8 |
| S2 | 165±33 | 48.9 |
| S3 | 43±15 | 12.7 |
| S4 | 46±29 | 13.6 |
| REM | 59±20 | 17.5 |
| DEL | 248±33 | - |

### MRI data acquisition and analysis

MR data acquisition took place at the Max-Planck-Institute for Biological Cybernetics, Tübingen, Germany in a 3T Magnetom Prisma scanner (Siemens, Erlangen, Germany) with a standard 20-channel head volume coil.

Functional images were obtained using multi-band sequences provided by the University of Minnesota Center for Magnetic Resonance Research. Functional data was acquired with an echo-planar sequence (cmrr_mbep2d_bold, 2300 ms repetition time [TR]; 30ms echo time [TE]; 75° flip angle [FA]; 208 mm field-of-view [FOV]; 104 voxel matrix size; 2.0×2.0×2.3 mm^3^ voxel size; 60 transversal slices with interleaved slice acquisition; parallel acquisition technique with inplane acceleration factor [GRAPPA] 2 and multi-band acceleration factor 2).

A high-resolution anatomical T1-weighted image was obtained during the second MR scanning block via a 3D magnetization-prepared 2 rapid gradient echoes (MP2RAGE) sequence (5000 ms TR; 2.98 ms TE; 700/2500 ms inversion time 1/2; 4°/5° FA 1/2; 256mm FOV; 256 voxel matrix size; 1 mm isotropic voxel size; 176 sagittal slices; GRAPPA 3). Also during the second block, a high-resolution T2-weighted image was acquired with a 3D turbo spin echo (TSE) sampling perfection with application optimized contrasts using different flip angle evolution (SPACE) sequence to improve pial surfaces within the FreeSurfer (v.5.3.0, http://surfer.nmr.mgh.harvard.edu) pipeline (3200 ms TR; 564 ms TE; 256 mm FOV; 320 voxel matrix size; 0.8 mm isotropic voxel size; 208 sagittal slices; GRAPPA 2). Slice acquisition occurred in an interleaved fashion. In addition to the functional and high-resolution anatomical data, a series of diffusion-weighted and control images were acquired, which were not analyzed for the current manuscript. Specifications are reported in Brodt et al. (2018).

Where applicable, data were preprocessed on the basis of the minimal preprocessing pipelines for the Human Connectome Project (Glasser et al., 2013) with the FMRIB Software Library (FSL, v6.0.1)(Jenkinson et al., 2012). Gradient nonlinearity distortion correction was omitted due to the lower gradient strength of 80mT/m compared to the HCP protocol. For data quality control, each correction or registration step was conducted separately and checked visually.

### Anatomical data

Initial brain extraction of the high-resolution anatomical T1 and T2-weighted scans was done via AC-PC alignment and subsequent linear (flirt) and nonlinear (fnirt) registration to the MNI space template, with the inverted warp bringing the mask back to native space. The T2 image was read-out distortion corrected and registered to T1 via epi_reg using the field map of the respective MR run. Both images were bias-field corrected, with the bias field estimated from the square root of the product of both images. Subsequently, transforms to MNI space were created via a 12 degrees of freedom (df) affine (flirt) and nonlinear (fnirt) registration. The corrected T1 and T2 images were fed into FreeSurfer’s recon -all pipeline, supporting brain extraction with the initially created brain mask and where required applying the skullstrip gcut option. The final brain mask used for the first-level functional data analyses was created based on FreeSurfer’s wmparc volume parcellation.

### Functional data

Preprocessing of the functional data followed the volumetric stream of the minimal preprocessing pipelines. The first five volumes of every run were discarded to control for magnetic saturation effects. The remaining volumes were motion corrected via a 6 df *mcflirt* registration to the single-band reference of the respective run. No data had to be excluded due to excessive movement (all translations <= 5 mm, rotations <= 5°). The single-band reference of each run was registered to each subject’s high -resolution T1 with *epi_reg* using the run’s respective B0 field map for distortion correction. Registration transforms and distortion corrections were concatenated to enable one single resampling step to 2 mm MNI space. Afterwards, data for each run was intensity-normalized to a brain mean of 10000, smoothed with a Gaussian kernel of 6 mm FWHM and high-pass filtered with a cutoff frequency of 0.01Hz.

Functional data was analyzed within a two-level model using SPM12 (Wellcome Department of Cognitive Neurology, London, UK). The first-level analysis was implemented on the subject-level modeling fixed effects. For the main model, conditions of interest were the encoding, retrieval and baseline trials separately for each encoding-retrieval run. Encoding trials were determined by the onset of the first item of a pair and the offset of the respective second item. Baseline trials lasted from the onset of the first fixation cross to the offset of the last fixation cross. The onset of retrieval trials was defined by the onset of the cue item, the trial offset was defined by the first button press, indicating that the participant had retrieved and chosen his answer location. This approach excluded potentially confounding activity due to the automatization of the finger-to-cursor-direction motor mapping. Changes in blood-oxygen level dependent (BOLD) responses were estimated by convolving the modeled data epochs with a canonical hemodynamic response function and estimating parameters using a restricted maximum likelihood approach at each voxel for every subject. The six head motion parameters from the motion correction step, as well as constants delineating each MR acquisition run were added as regressors of no interest.

The first second-level random effects model was a full factorial analysis with the within-subject factor repetition and the between-subject factor group (wake/sleep) on the original, unadjusted data (see below). It was based on first-level contrast images that subtracted the respective baseline activation from the condition of interest and were spatially smoothed with a Gaussian kernel of 8 mm FWHM.

For a conservative analysis that excludes any potential effects of time spent awake at the time of data recording, we estimated an additional second-level model including all retrieval repetitions as a single condition of interest and a covariate modeling the time spent awake for each subject and each task session (sleep group S1: 750±74 min [M±SD]; sleep group S2: 160±27 min [M±SD]; wake group S1: 169±39 min [M±SD]; wake group S2: 969±46 min [M±SD]). The residuals of this model were then subjected to a further second-level model analogous to the main model, enabling the detection of sleep effects now free of any effects of time spent awake. Regions of interest (ROIs) for further independent analyses were determined based on significant clusters in this analysis in regions known for their involvement in memory-related processing.

Like with the behavioral data, we assessed sleep effects on functional brain activity with an interaction contrast between the two groups and retrieval activity in the last run of the first task session (‘pre’) and the first run of the second task session (‘post’).

A psychophysiological interaction (PPI) analysis was performed using an anatomical mask of the ROIs showing significant sleep effects as seed. For assessing sleep-specific changes in connectivity, estimation of the single-subject GLM was based on the two task runs directly before and after the retention interval (pre/post). The model included 9 regressors per run: the deconvolved first eigenvariate extracted from the signal of the seed region, the trial onset vector for the single run convolved with the HRF, the interaction term of the eigenvariate with the onset vector and the six head motion parameters. As with the behavioral and functional activity analysis, sleep effects were assessed by testing the interaction between group (wake/sleep) and time point (pre/post).

A custom-built MATLAB script was used for the correlation analysis between functional data and behavioral performance, which correlated the average functional parameter estimates across all voxels in the respective ROI with the behavioral data across participants. As above, retrieval performance was correlated with the difference in retrieval -related functional activity between pre and post retention runs. Because of outliers in the functional parameter estimates (>M±3SD) we report Spearman correlations.

If not indicated otherwise, statistical inferences on whole-brain functional data were obtained at a threshold of *p*_uncorr_<.001 and then verified with small-volume FWE correction *p*_SVC_<.05 for the anatomical region of the respective significant cluster (Harvard-Oxford cortical/subcortical structural atlas, thresholded at minimum probability of 25% as included in FSL v.6.0.1). To assess the direction of the interaction effects in the ROIs, post-hoc tests were performed on the mean beta values of all significant voxels of activated clusters as well as on the mean beta values of all voxels within the corresponding anatomical regions with repeated measures ANOVAs and two-sided t-tests.

In general, significant clusters reported in the text are provided with their peak voxel MNI-coordinates and corresponding t-values in the supplementary tables, together with all significant clusters with an extent above 10 voxels from the whole-brain analyses. Results are displayed on MRIcroGL’s MNI152_2009 template (University of South Carolina, Columbia, SC). For all activated clusters, anatomical localization was determined based on automatic labeling of the largest region within a cluster with FSL’s autoaq based on the Harvard-Oxford cortical/subcortical structural atlas labels.

## Author contributions

S.B., M.S., M.E., K.S., and S.G. designed the research; S.B., M.S., A.S., J.B., and M.E. performed the experiments; S.B., A.S., and J.B. analyzed the functional and behavioral data; S.B., M.S. and S.G. wrote the manuscript.

