## Supplementary material for "Memory systems integration in sleep complements rapid systems consolidation in wakefulness": SI

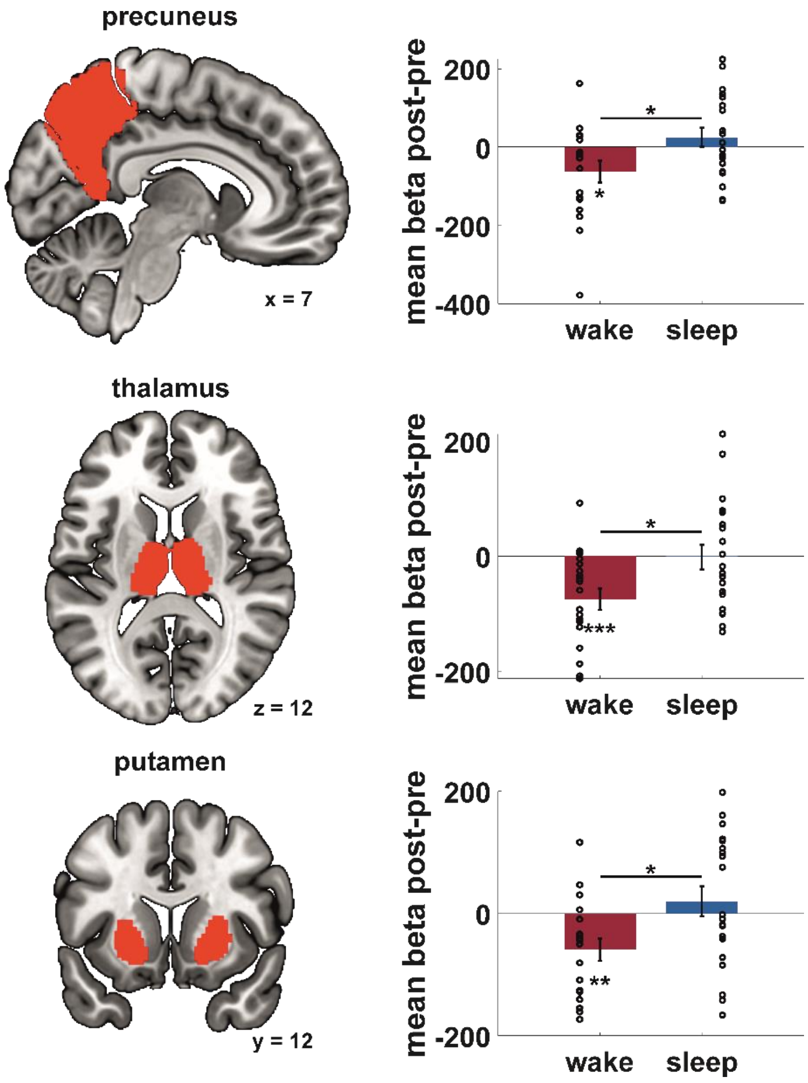


**Fig. S1.**

**Sleep-dependent increases in retrieval activity across the anatomically defined ROIs from the significant clusters in Fig. 3.** (**A**) The mean beta values across the anatomically defined precuneus, thalamus and putamen voxels show a significant modulation of retrieval activity by sleep (interaction group*time). Red, wake; blue, sleep; dots represent individual data points. See Table S3C. Across the whole anatomical regions, the interaction is driven by a concurrent decrease in retrieval activity in the wake group and stabilization in the sleep group. All beta values are corrected for baseline activation levels. Data are M±SEM. * $p<.05$, ** $p<.01$, *** $p<.001$.

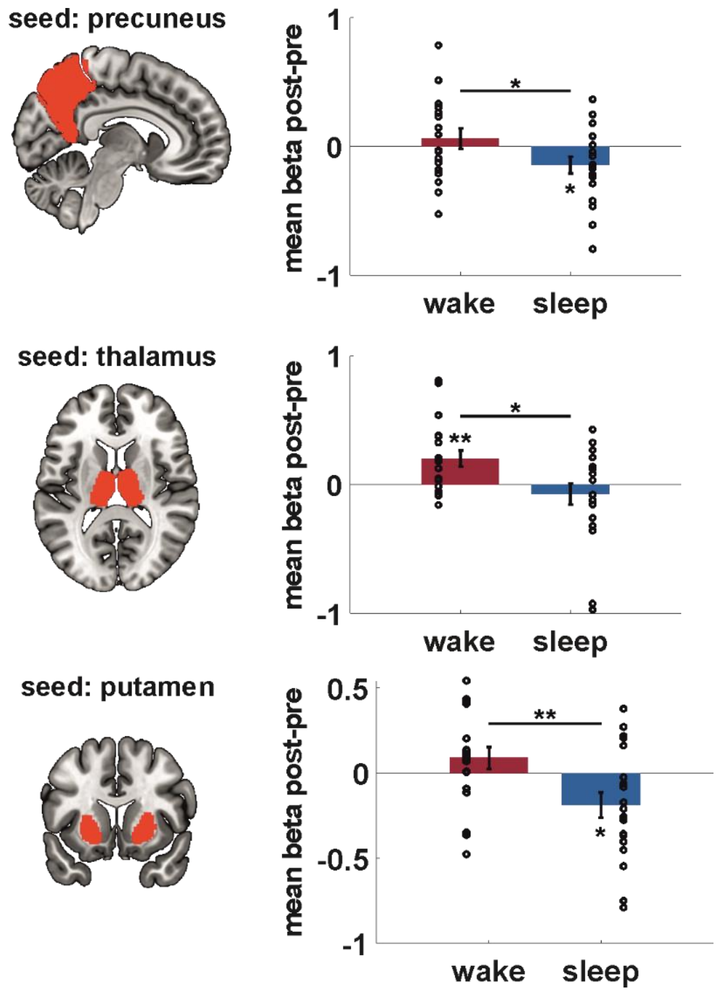


**Fig. S2.**

**Sleep-dependent decrease in memory-related functional connectivity across the whole anatomically defined hippocampus ROI.** C.f. Fig 5B. Complementing the main analysis, investigation of the mean beta values of the anatomical hippocampus voxels shows an interaction effect with differential changes in connectivity in the wake and sleep groups across the retention interval. Concurrent increase in connectivity with the hippocampus in the wake and decrease in the sleep group across the retention interval (Table S6B). Data are M±SEM. Dots represent individual data points. All beta values are corrected for baseline activation levels. * $p<.05$, ** $p<.01$.

**Table S1.**

**Sleep effects on memory-related functional activity controlled for influence of sleep pressure.** All regions listed exhibited significant peak voxel effects at $p_{uncorr}<.001$, df=296 and had a cluster size≥10 voxels. *Regions also showed significant peak-voxel-effects at $p_{SVC}<.05$ in an analysis of the anatomically defined ROI. Clusters also did not show a significant time-of-day effect ($T_{tod}$). Peak voxel MNI coordinates are given in mm. Assigned region is the region from the Harvard-Oxford Cortical/Subcortical Atlas constituting the largest portion of the cluster.

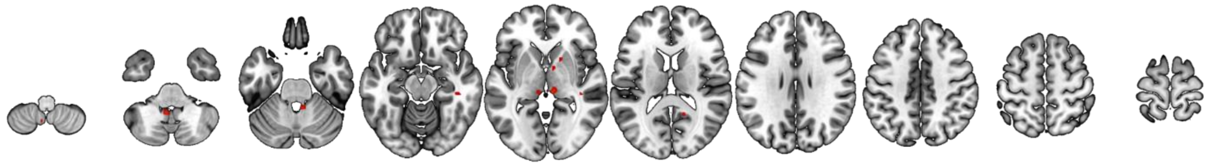

| Region | n voxels | $T_{peak}$ | $x_{peak}$ | $y_{peak}$ | $z_{peak}$ | $T_{tod}$ |
|---|---|---|---|---|---|---|
| Cerebellum | 72 | 3.71 | -6 | -52 | -40 | 1.16 |
| Thalamus* | 63 | 3.66 | 6 | -28 | -2 | -0.16 |
| Precuneus* | 33 | 3.34 | 14 | -54 | 18 | 0.44 |
| Putamen* | 29 | 3.29 | 8 | 0 | -2 | 1.81 |
| Fusiform Cortex | 24 | 3.31 | 44 | -30 | -10 | 1.76 |
| Cerebellum | 21 | 3.39 | 6 | -44 | -30 | -0.05 |
| Cerebellum | 19 | 3.46 | -4 | -60 | -54 | -0.17 |
| Thalamus* | 16 | 3.39 | -8 | -28 | 0 | 0.09 |
| Putamen* | 14 | 3.21 | -20 | 10 | -6 | -0.16 |
| Middle Temporal Gyrus | 14 | 3.26 | 34 | 6 | 48 | 0.73 |

**Table S2.**

**Regions displaying significant sleep effects in functional activity**. All regions listed exhibited significant peak voxel effects at $p_{uncorr}$ <.001, df=296 and had a cluster size≥10 voxels. *Regions also showed significant peak-voxel-effects at $p_{SVC}$<.05 in an analysis of the anatomically defined respective ROI. Peak voxel MNI coordinates are given in mm. Assigned region is the region from the Harvard-Oxford Cortical/Subcortical Atlas constituting the largest portion of the cluster.

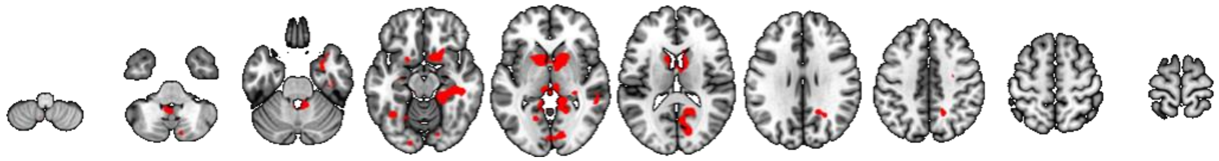

| Region | n voxels | $T_{peak}$ | $x_{peak}$ | $y_{peak}$ | $z_{peak}$ |
|---|---|---|---|---|---|
| | 2716 | 4.36 | 2 | -30 | -2 |
| Intracalcarine Cortex | 271 | 3.94 | 18 | -50 | 8 |
| incl. Precuneus* | 48 | 4.01 | -6 | -30 | 0 |
| incl. Thalamus* | 47 | 3.85 | 6 | -26 | 0 |
| Putamen* | 1949 | 4.55 | 20 | 12 | -6 |
| Cerebellum | 362 | 4.31 | -4 | -50 | -38 |
| Occipital Pole | 90 | 3.76 | 28 | -98 | -10 |
| Precentral | 78 | 3.54 | 44 | 0 | 50 |
| Middle Temporal Gyrus | 65 | 3.36 | 56 | -42 | 0 |
| Temporal Pole | 52 | 3.54 | 30 | 4 | -28 |
| Fusiform Gyrus | 46 | 3.48 | -38 | -58 | -14 |
| Occipital Pole | 35 | 3.28 | -16 | -92 | -14 |
| Lingual Gyrus | 35 | 3.41 | -18 | -62 | 0 |
| Hippocampus | 31 | 3..46 | -34 | -14 | -20 |
| Fusiform Gyrus | 30 | 3.26 | 16 | -80 | -16 |
| Precentral | 29 | 3.42 | 28 | -10 | 46 |
| Cerebellum | 14 | 3.29 | 12 | -80 | -44 |
| Inferior Frontal Gyrus | 13 | 3.34 | -60 | 28 | 20 |
| Fusiform Gyrus | 10 | 3.2 | -22 | -62 | -14 |
| Parahippocampal Gyrus | 10 | 3.32 | -14 | -32 | -14 |

**Table S3.**

**Sleep effects on memory-related functional activity**. Statistics on the mean beta values for the precuneus, thalamus and putamen clusters display sleep-related changes in functional activity during retrieval across time points. Repeated-measures ANOVAs with the between factor group (sleep vs. wake) and within factor time (post vs. pre), n=39. F-, p- and partial η2-values of the interaction effect, t- and p-values of the t-tests comparing time points within the wake (W) and sleep (S) groups.

| Region | $F_{1,37}$ | p | $\eta^2$ | $t_{W(18)}$ | $p_W$ | $t_{S(19)}$ | $p_S$ |
|---|---|---|---|---|---|---|---|
| **A** – significant voxels s-w*post-pre residuals data | | | | | | | |
| Precuneus | 8.64 | .006 | .189 | 1.97 | .065 | -2.19 | .041 |
| Thalamus | 26.29 | <.001 | .415 | 5.23 | <.001 | -2.19 | .041 |
| Putamen | 34.56 | <.001 | .483 | .540 | <.001 | -3.12 | .006 |
| **B** – significant voxels s-w*post-pre original data | | | | | | | |
| Precuneus+Thalamus cluster | 30.94 | <.001 | .455 | 5.32 | <.001 | -2.66 | .015 |
| Putamen cluster | 23.83 | <.001 | .392 | 4.55 | <.001 | -2.25 | .036 |
| **C** – anatomically defined ROIs s-w*post-pre | | | | | | | |
| Precuneus | 5.75 | .022 | .135 | 2.30 | .034 | -1.01 | .325 |
| Thalamus | 6.71 | .014 | .154 | 3.29 | .004 | -0.82 | .423 |
| Putamen | 6.55 | .015 | .150 | 3.99 | <.001 | .04 | .966 |

**Table S4.**

**Regions displaying significant time-of-day effects in functional activity**. All regions listed exhibited significant peak voxel effects at $p_{uncorr}$ <.001, df=295 and had a cluster size≥10 voxels. Peak voxel MNI coordinates are given in mm. Assigned region is the region from the Harvard-Oxford Cortical/Subcortical Atlas constituting the largest portion of the cluster.

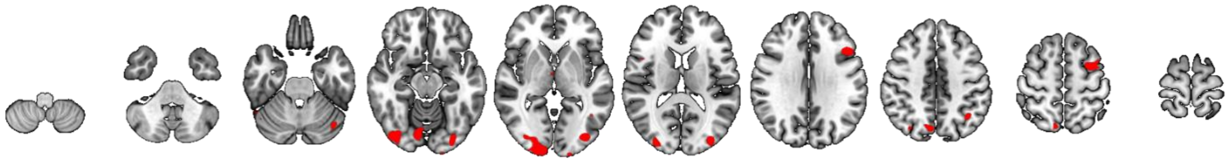

| Region | n voxels | $T_{peak}$ | $x_{peak}$ | $y_{peak}$ | $z_{peak}$ |
|---|---|---|---|---|---|
| Occipital Pole | 1193 | 4.94 | 10 | -104 | -2 |
| Lateral Occipital Cortex | 529 | 4.45 | -36 | -84 | -4 |
| Middle Frontal Gyrus | 212 | 3.89 | -46 | 18 | 30 |
| Middle Frontal Gyrus | 194 | 3.92 | -42 | 6 | 56 |
| Precuneus | 173 | 4.3 | 8 | -74 | 50 |
| Occipital Pole | 101 | 3.95 | -20 | -106 | -10 |
| Angular Gyrus | 96 | 4.09 | -38 | -54 | 38 |
| Cerebellum | 71 | 3.52 | -38 | -76 | -24 |
| Occipital Pole | 50 | 3.54 | 30 | -90 | 14 |
| Right Thalamus | 39 | 3.66 | 2 | -6 | -4 |
| Cerebellum | 38 | 3.55 | -6 | -74 | -24 |
| Inferior Frontal Gyrus | 32 | 3.41 | 46 | 10 | 18 |
| Lateral Occipital Cortex | 22 | 3.48 | 32 | -74 | 48 |
| Cerebellum | 11 | 3.34 | 38 | -66 | -52 |

**Table S5.**

**Regions displaying significant sleep effects on functional connectivity**. All regions listed exhibited significant peak voxel effects at $p_{uncorr}<.001$, df=296 and had a cluster size≥10 voxels. *Regions also showed significant peak-voxel-effects at $p_{SVC}<.05$ in an analysis of the anatomically defined respective ROI. Peak voxel MNI coordinates are given in mm. Assigned region is the region from the Harvard-Oxford Cortical/Subcortical Atlas constituting the largest portion of the cluster.

| Region | n voxels | $T_{peak}$ | $x_{peak}$ | $y_{peak}$ | $z_{peak}$ |
|---|---|---|---|---|---|
| **A** – PPI seed: precuneus | | | | | |
| 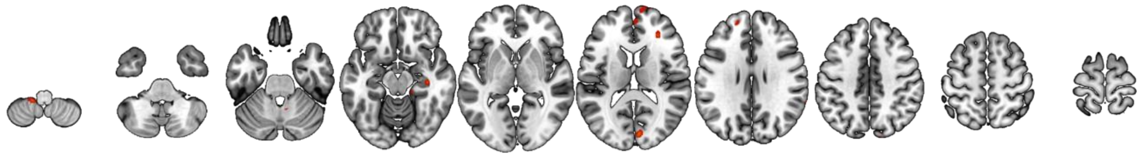 | | | | | |
| Occipital Pole | 228 | 4.17 | 6 | -92 | 20 |
| Paracingulate Cortex | 122 | 3.68 | -8 | 48 | 8 |
| Cerebellum | 66 | 4.05 | 10 | -56 | -36 |
| Precentral Gyrus | 66 | 3.83 | 8 | -22 | 64 |
| Frontal Pole | 36 | 3.39 | 10 | 68 | 14 |
| Paracingulate Gyrus | 36 | 3.56 | -6 | 38 | 34 |
| Cerebellum | 31 | 3.61 | -14 | -44 | -58 |
| Parahippocampal Gyrus | 30 | 3.59 | 18 | -34 | -10 |
| Frontal Pole | 23 | 3.64 | 30 | 36 | 14 |
| Hippocampus* | 21 | 3.65 | 36 | -22 | -14 |
| Angular Gyrus | 16 | 3.56 | 68 | -46 | 26 |
| Frontal Pole | 16 | 3.36 | -18 | 50 | 30 |
| Lingual Gyrus | 15 | 3.5 | -2 | -66 | -4 |
| **B** – PPI seed: thalamus | | | | | |
| 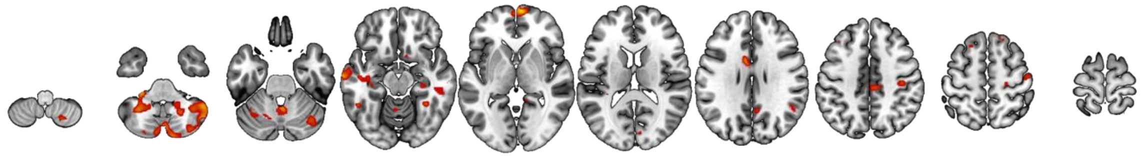 | | | | | |
| Cerebellum | 1306 | 4.31 | -18 | -58 | -22 |
| Cerebellum | 1006 | 4.39 | 52 | -54 | -40 |
| Middle Temporal Gyrus | 564 | 4.32 | -68 | -16 | -10 |
| Cerebellum | 477 | 3.74 | 4 | -76 | -42 |
| Cingulate Gyrus | 354 | 4.38 | 24 | -24 | 50 |
| Angular Gyrus | 301 | 3.92 | 48 | -54 | 34 |

| | | | | | |
|---|---|---|---|---|---|
| Frontal Pole | 294 | 4.43 | 6 | 66 | -2 |
| Middle Temporal Gyrus | 280 | 4.12 | 50 | -30 | -10 |
| Cingulate Gyrus | 265 | 4.31 | -6 | 2 | 32 |
| Precuneus | 115 | 3.53 | 6 | -58 | 34 |
| Postcentral Gyrus | 68 | 3.41 | 50 | -14 | 56 |
| Hippocampus* | 65 | 3.45 | 32 | -26 | -10 |
| Fusiform Cortex | 47 | 3.59 | 34 | -48 | -16 |
| Frontal Pole | 39 | 3.94 | 12 | 40 | 58 |
| Cuneal Cortex | 33 | 3.4 | 6 | -84 | 18 |
| Intracalcarine Cortex | 31 | 3.36 | 20 | -80 | 6 |
| Cerebellum | 30 | 3.53 | 20 | -66 | -58 |
| Fusiform Cortex | 25 | 3.44 | -36 | -50 | -22 |
| Cerebellum | 25 | 3.47 | -54 | -54 | -50 |
| Inferior Temporal Gyrus | 24 | 3.49 | -50 | -50 | -16 |
| Cerebellum | 20 | 3.29 | -24 | -84 | -42 |
| Planum Temporale | 18 | 3.31 | -32 | -34 | 12 |
| Subcallosal Cortex | 17 | 3.5 | 10 | 12 | -18 |
| Frontal Orbital Cortex | 16 | 3.57 | 38 | 30 | -22 |
| Parahippocampal Gyrus | 14 | 3.35 | 22 | -34 | -12 |
| Cingulate Gyrus | 11 | 3.3 | 12 | -44 | 2 |
| Middle Frontal Gyrus | 11 | 3.28 | -38 | 30 | 44 |
| Superior Frontal G | 10 | 3.25 | -20 | 22 | 58 |

**C** – PPI seed: putamen

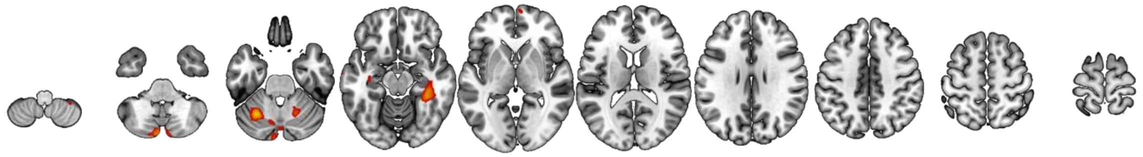

| | | | | | |
|---|---|---|---|---|---|
| Cerebellum | 476 | 4.46 | -6 | -88 | -38 |
| Cerebellum | 386 | 4.3 | -34 | -58 | -32 |
| Fusiform Cortex | 277 | 3.99 | 36 | -38 | -14 |
| Middle Frontal Gyrus | 119 | 3.75 | 36 | 26 | 52 |
| Cerebellum | 74 | 3.59 | 20 | -58 | -28 |
| Putamen | 69 | 3.58 | -32 | -16 | -8 |
| Precuneus | 56 | 3.37 | 8 | -58 | 18 |
| Cerebellum | 53 | 3.97 | -54 | -54 | -50 |
| Frontal Pole | 38 | 3.39 | 6 | 66 | 2 |
| Precuneus | 33 | 3.36 | -10 | -64 | 20 |
| Frontal Orbital Cortex | 33 | 3.46 | 30 | 26 | -22 |
| Frontal Orbital Cortex | 24 | 3.41 | -22 | 28 | -20 |
| Middle Temporal Gyrus | 15 | 3.29 | -70 | -10 | -16 |
| Lateral Occipital Cortex | 12 | 3.27 | 40 | -58 | 40 |
| Cerebellum | 11 | 3.36 | 56 | -54 | -34 |
| Hippocampus* | 10 | 3.4 | 18 | -10 | -26 |

### Table S6

**Sleep effects on memory-related functional connectivity with the hippocampus**. Statistics on the mean beta values display sleep-related changes in functional connectivity between the precuneus, putamen and thalamus, respectively, and the hippocampus during retrieval across time points. Repeated-measures ANOVAs with the between factor group (sleep vs. wake) and within factor time (post vs. pre), n=39. F-, p- and partial η2-values of the interaction effect, t- and p-values of the t-tests comparing time points within the wake (W) and sleep (S) groups.

| Region | $F_{1,37}$ | p | $\eta^2$ | $t_{W(18)}$ | $p_W$ | $t_{S(19)}$ | $p_S$ |
|---|---|---|---|---|---|---|---|
| **A** – significant hippocampus clusters | | | | | | | |
| Precuneus | 20.03 | <.001 | .351 | -2.57 | .019 | 4.13 | <.001 |
| Thalamus | 23.89 | <.001 | .392 | -4.45 | <.001 | 2.96 | .008 |
| Putamen | 18.05 | <.001 | .328 | -2.46 | .024 | 3.52 | .002 |
| **B** – anatomically defined hippocampus ROI | | | | | | | |
| Precuneus | 4.20 | .047 | .102 | -0.80 | .435 | 2.24 | .037 |
| Thalamus | 6.79 | .013 | .155 | -3.18 | .005 | 0.87 | .396 |
| Putamen | 7.93 | .008 | .176 | -1.36 | .192 | 2.58 | .018 |
